# DAFi: A Directed Recursive Filtering and Clustering Approach to Data-Driven Identification of Cell Populations from Polychromatic Flow Cytometry Data

**DOI:** 10.1101/193912

**Authors:** Alexandra J. Lee, Ivan Chang, Julie G. Burel, Cecilia S. Lindestam Arlehamn, Daniela Weiskopf, Bjoern Peters, Alessandro Sette, Richard H. Scheuermann, Yu Qian

## Abstract

Computational methods for identification of cell populations from high-dimensional flow cytometry data are changing the paradigm of cytometry bioinformatics. Data clustering is the most common computational approach to unsupervised identification of cell populations from multidimensional cytometry data. We found that combining recursive filtering and clustering with constraints converted from the user manual gating strategy can effectively identify overlapping and rare cell populations from smeared data that would have been difficult to resolve by either a single run of data clustering or manual segregation. We named this new method DAFi: Directed Automated Filtering and Identification of cell populations. Design of DAFi preserves the data-driven characteristics of unsupervised clustering for identifying novel cell-based biomarkers, but also makes the results interpretable to experimental scientists as in supervised classification through mapping and merging the high-dimensional data clusters into the user-defined 2D gating hierarchy. By recursive data filtering before clustering, DAFi can uncover small local clusters which are otherwise difficult to identify due to the statistical interference of the irrelevant major clusters. Quantitative assessment of cell type specific characteristics demonstrates that the population proportions calculated by DAFi, while being highly consistent with those by expert centralized manual gating, have smaller technical variance than those from individual manual gating analysis. Visual examination of the dot plots showed that the boundaries of the DAFi-identified cell populations followed the natural shapes of the data distributions. To further exemplify the utility of DAFi, we show that DAFi can incorporate the FLOCK clustering method to identify novel cell-based biomarkers. Implementation of DAFi supports options including clustering, bisecting, slope-based gating, and reversed filtering to meet various auto-gating needs from different scientific use cases.

## 1. Introduction

The success of flow cytometry (FCM) is dependent on being able to accurately identify discriminant cell populations. Currently, the most common existing approach is manual gating analysis. In a typical manual gating procedure, an experiment operator would start by inspecting the distribution of cellular events on a selected pair of measured characteristics (scatter parameters or protein markers) on the 2D plot, visually recognize the clusters of the cellular events, draw a 2D polygon to extract a population of interest, inspect the population on another pair of markers to identify its subpopulations, and repeat this procedure to further partition each subpopulation until all cell subsets of interest are identified. Through this 2D by 2D recursive segregation, cell populations are identified and managed in a user-defined hierarchy with phenotypes defined with the markers used at each gating step. Desirable features of manual gating analysis include the flexibility in the analysis procedure and the interpretability of the analysis results.

However, the manual gating procedure is also subjective, time-consuming, and difficult to reproduce. Technical variance is usually found in independent manual gating analysis conducted across experiments, studies, and labs [12, 31]. Except in the ideal case where each cell population is highly cohesive and segregated from others, there is notable bias in using a sequential 2D by 2D analysis to identify cell populations defined in high-dimensional space since cell populations can be difficult to separate on 2D dot plots. Manual gating analysis typically bisects the overlapped populations with manually drawn lines, resulting in inaccurate identification and calculation of the population characteristics. Additionally, the design of the manual gating approach is not suitable for exploratory data analysis. Gating steps are predefined on user-selected markers and constrained by the operator’s knowledge of cell population phenotypes. Recent advance in cytometry instrumentation and reagent technology made these issues more severe (*e.g.*, Becton Dickinson’s FACSymphony™ is claimed to be able to measure up to 50 different characteristics). With these many parameters, it becomes almost impossible to explore the enormous high-dimensional data space exhaustively and accurately using a manual gating approach, considering the time, effort, and human bias involved in the analysis.

During the last decade, many computational methods have been developed for the identification of cell populations from polychromatic FCM data. State-of-the-art computational approaches are shown to be superior to manual gating analysis in terms of efficiency, reproducibility and reduction of human bias [2, 3, 8, 9, 10, 13, 21, 22, 25, 26, 38]. Based on whether user inputs are required, these approaches can be broadly categorized into unsupervised [1, 11, 14, 15, 18, 24, 28, 32, 33, 44, 46] and supervised/semi-supervised [16, 23, 27] approaches. Unsupervised methods are usually based on data clustering methods, making them useful for comprehensive immunophenotyping and identification of novel cell subsets. Because this unsupervised analysis is completely data-driven, there is no direct connection between the identified data clusters and existing knowledge about the cell populations. Each data cluster needs to be annotated and validated, usually manually. The number of clusters identified in different input files can also be different. Therefore, it can be non-trivial to map and interpret all cell populations identified by the unsupervised clustering methods across samples. In contrast, supervised identification methods require prior data analysis results (usually from manual gating analysis) as training data, and thereby guarantee the interpretability of the identified data clusters. The trade-off is that the supervised methods, primarily focused on predefined cell populations, usually do not support exploratory discovery of novel cell subsets. While identifying novel cell subsets is one of the most important features expected by translational researchers when they use computational methods for FCM data analysis, supervised identification methods are preferred for clinical diagnostics.

Because there is no single model that fits all data, the biggest challenge for adoption of a computational method by experimental scientist is how to select the best method when the dataset changes. Parameters in a computational algorithm often need to be adjusted when being applied to a new dataset, and the adjusting is usually difficult without sufficient understanding about the method and the data. This challenge may be partially addressed with infrastructure efforts such as developing a parallel testing environment (*e.g.*, FlowGate [35]) to assess the performance of multiple applicable computational methods on each specific dataset. Another solution is to support the incorporation of user knowledge to guide the clustering analysis. One example is constrained clustering, in which user-provided constraints about cluster membership of data objects are involved [45]. Constrained clustering is regarded as a special class of semi-supervised learning, which has proven highly effective for solving domain-specific problems.

In this paper, we propose a constraint-based recursive filtering and clustering approach – DAFi (directed automated filtering and identification of cell populations) – to address the problem of utilizing computational methods for identification of cell populations from FCM data. Our goal is not to propose a new data clustering or classification method. Instead, we will demonstrate that designing a recursive filtering and clustering approach and combining it with user gating strategy can effectively and reliably accomplish the task of auto-gating for identifying not only the major but also rare and novel cell populations from a variety of FCM datasets.

## 2. Results

Figure 1 illustrates the design of DAFi. At each gating step DAFi supports the use of different clustering methods as well as bisecting (for identifying outlier cells), slope-based (e.g., identifying singlets using FSC/SSC-A vs FSC/SSC-H), and reverse-gating (events inside the hyper-polygon will be filtered out). We experimented with two clustering methods: *K*-means and FLOCK clustering [33], for benchmarking the performance of DAFi. User input is used to identify both predefined and novel populations that can be organized within an easily interpreted gating hierarchy (Figure 1B). We refer to this type of approach as *directed unsupervised clustering*. Figure 1C shows DAFi-identified major (CD4+ T and CD8+ T cells) and rare (CD3+CD56+ T and CD3hiCD56+ T cells) cell populations. The CD3+CD56+ T and CD3hiCD56+ T cell populations are difficult to separate by either manual gating or traditional unsupervised data clustering methods because the two clusters are both relatively rare and close to each other in CD3 expression distributions. However, they were well segregated with natural boundaries (unimodal distribution on each dimension) using DAFi, which applied recursive clustering with manual gating polygons as *constraints* rather than absolute boundaries. DAFi also identified the difficult-to- resolve CD4+CD25+ regulatory T cells (Tregs), yielding cell populations with natural distributions (Figure 1D). In contrast, manual gating analysis using polygon partitions did not capture natural boundaries of cell populations (*i.e.*, an abrupt lower boundary in the CD25 dimension); *K*-means clustering failed to identify the rare Treg cell population at all, even with *K=500*.

**Figure 1.**
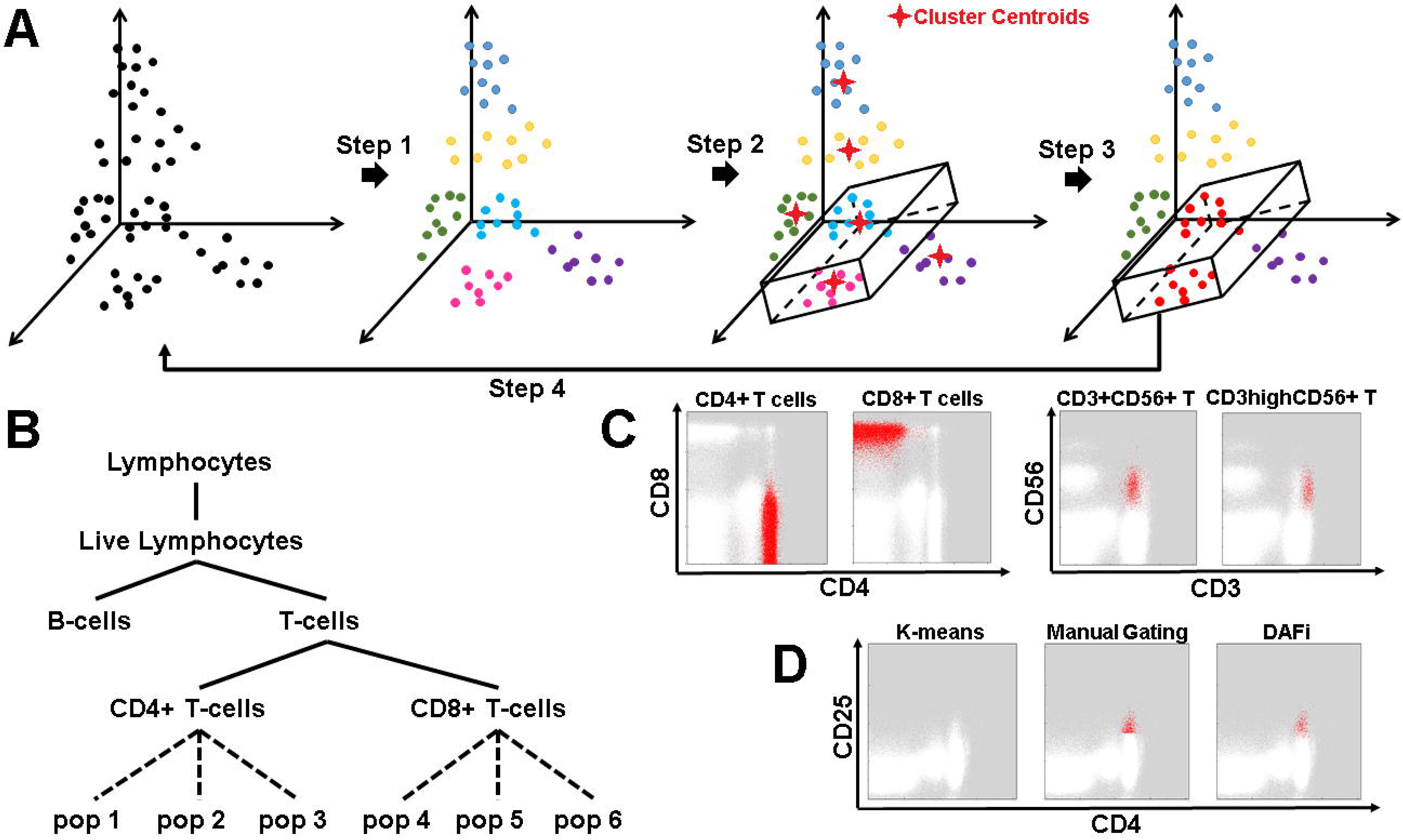
Design features of DAFi. A) Steps in the DAFi workflow. In Step 1, putative cell populations are identified by data clustering in high dimensional space, with cell events colored by population membership. In Step 2, a hyper-polygon is provided from combining 2D manual gating boundaries to identify the dataspace region of interest. Cell clusters are selected if their centroids are located within the hyper-polygon (two clusters shown, in light blue and magenta). In Step 3, all cell events associated with the centroids are selected and retained as the filtered population (in red), which is used as the input to the next iteration in Step 4. B) An example gating hierarchy in which the DAFi framework can be used to identify both predefined (solid lines) and novel (dotted lines) cell populations, and organize them within a user-provided gating hierarchy for simplified annotation and interpretation. C) DAFi identification of CD4+T, CD8+T, CD3+CD56+ T and CD3hiCD56+ T cells. CD4+T and CD8+T cells are shown on CD4 vs CD8 dot plots, while CD3+CD56+ T and CD3hiCD56+ T cells are on CD3 vs CD56 plots. Cell populations identified by DAFi are colored in red. D) Comparison of DAFi with other clustering and filtering methods. Putative CD4+CD25+ regulatory T cells (Tregs) were identified using *K*-means clustering, manual gating, and DAFi. The identified Treg cells are colored in red and the remaining cells colored in white.

We evaluated the performance of DAFi using FCM data from both the public ImmPort database (Immunology Database and Analysis Portal, http://www.immport.org) and our HIPC (Human Immunology Project consortium, https://www.immuneprofiling.org) studies. Results of DAFi were assessed both quantitatively and by visual examination of the identified cell populations on dot plots. For the quantitative assessment, instead of using the “bulk assessment” such as the sample-level F-measure which is dominated by contributions from the abundant cell populations, we focus on cell type specific statistics for each individual cell population.

### 2.1 Cell Type Specific Assessment in Comparison with Individual and Centralized Manual Gating Analysis

The first assessment focused on the identification of different T cell subsets using a representative 10- color reagent panel on multiple repeat runs of cryopreserved PBMC (peripheral blood mononuclear cells) from one sample donation of a healthy donor [12]. Repeated FCM experiments were performed on various days throughout a 7-month period by three different operators on four different cytometers. Technical variability associated with each cell population across the 24 runs can therefore be estimated and compared between the results of DAFi and those from the manual gating analysis.

Two manual gating analysis results were available for comparison - individual manual gating analysis (INDI) performed by different operators when the FCM data were acquired, and centralized manual gating analysis (CENT) performed by one analyst after data from all 24 samples had been acquired. The variety of cell subsets and their relationship specified by expert manual gating in a predefined hierarchy are shown in Figure 2A (also in Supplementary File 1). Among the 22 cell populations identified by manual gating analysis, 17 of them are of special user interest. They were divided into two categories based on the technical variance – “clearly defined” and “poorly resolved” - defined by coefficient of variability (CV) in cell population proportions across the 24 samples from individual manual gating analysis:

- Clearly defined (low CV): 5: Monocytes; 7: B-cells; 8: NK cells; 11: T-cells; 12: CD4+ T cells; 13: CD8+ T cells; 15: Naïve CD4+ T cells; 19: Naïve CD8+ T cells.
- Poorly resolved (high CV): 9: CD3+CD56+ T cells; 10: CD3highCD56+ T cells; 14: Tregs (regulatory CD4+ T cells); 16: Tcm CD4+ T cells (central memory CD4+ T cells); 17: Tem CD4+ T cells (effector memory CD4+ T cells); 18: Temra CD4+ T cells (effector memory CD4+ T cells that express CD45RA); 20: Tcm CD8+ T cells; 21: Tem CD8+ T cells; 22: Temra CD8+ T cells.

**Figure 2.**
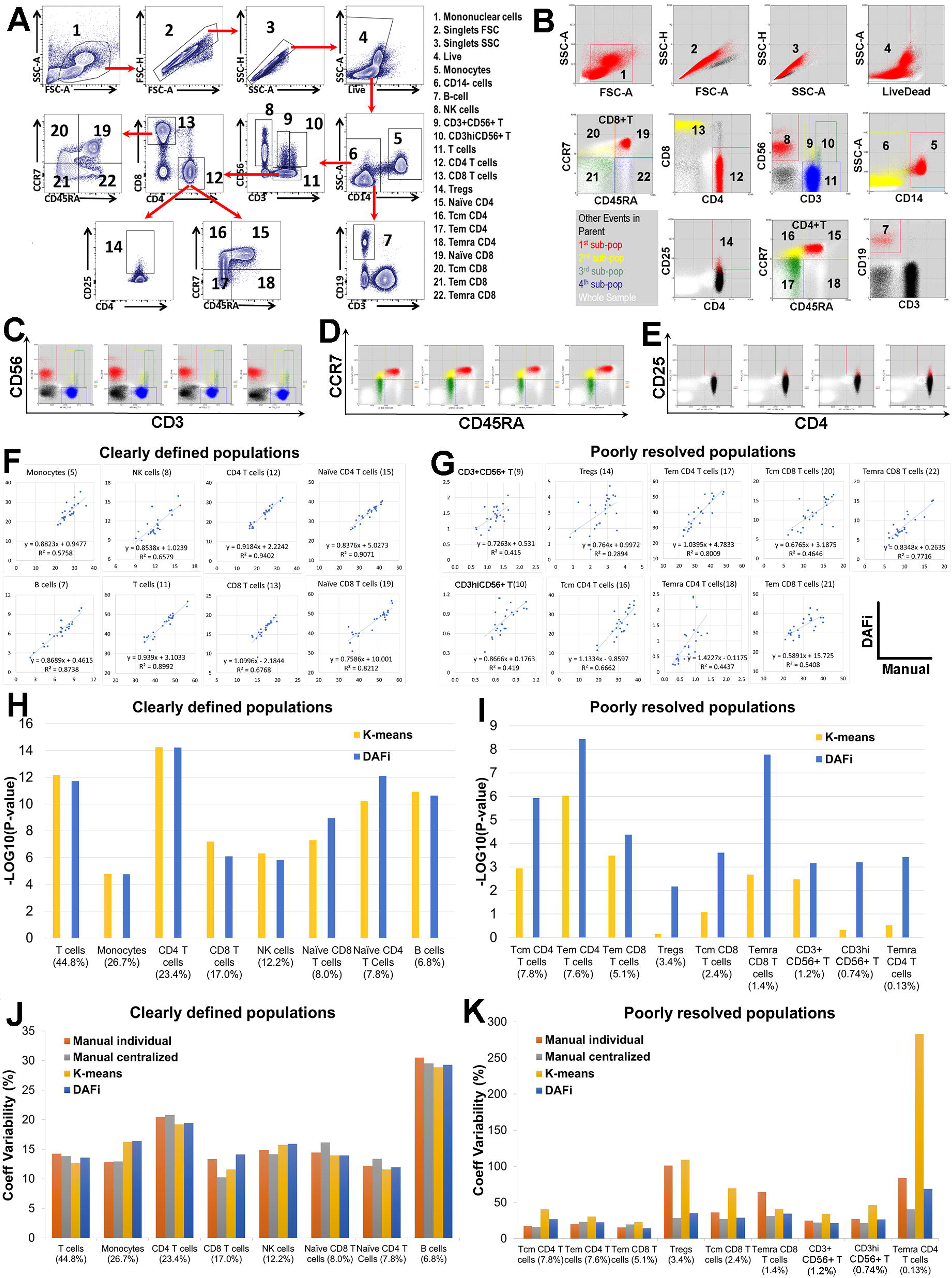
Performance evaluation of DAFi in comparison with individual and centralized manual gating analysis. A) Illustration of the manual gating hierarchy for identifying the 22 predefined cell populations from the 10-color T cell panel, with gating boundaries shown on each 2D dot plot. Along the direction of the red arrows is the sequence of the gates with their parent populations. The cell populations are numbered. Names of the cell types are listed to the right. B) DAFi results for identifying the corresponding 22 predefined cell populations. Events from the whole sample are colored in white. The black colored dots are events of the parent population, with events identified by DAFi highlighted in red, yellow, green, and blue. C) T cells (Blue), NK cells (Red), CD3+CD56+ T (yellow), and CD3hiCD56+ T (green) from four data files are shown. D) Naïve CD4+ T (red), effector memory CD4+ T (Tem CD4, green), central memory CD4+ T (Tcm CD4, yellow), and effector memory CD4+ T expressing CD45RA (Temra CD4, blue) identified by DAFi from four data files are shown. E) Regulatory T cells (Tregs, CD4+CD25+, red) identified by DAFi from four data files are shown. F) Linear regression and correlation analysis of cell population percentages of clearly-defined cell populations identified by DAFi (*y*-axis) compared with centralized manual gating (*x*-axis). G) Linear regression and correlation analysis of percentages of poorly-resolved cell populations identified by DAFi (*y*-axis) compared with centralized manual gating analysis (*x*-axis). H) P-values (-log10 transformed) of *x*-variable in linear regression analysis between percentages generated by data clustering methods (DAFi and *K*-means) and centralized manual gating analysis for clearly-defined cell populations. The cell populations were sorted based on their average percentage of their parents from the largest to the smallest, as shown on the *x*-axis. I) P-values (-log10 transformed) of linear regression analysis between percentages generated by data clustering methods (DAFi and *K*-means) and centralized manual gating analysis for poorly-resolved cell populations. J) Coefficient variability (CV) of population percentages across the 24 samples for clearly- defined cell populations. K) CV of population percentages across the 24 samples for poorly-resolved cell populations.

Previously [12], we found that both INDI and CENT could achieve a high degree of concordance for identifying clearly defined cell populations; but for the poorly resolved ones, CENT significantly outperformed INDI.

Figure 2B shows the results of DAFi. Visual examination shows the main difference between DAFi and the manual gating analysis is that DAFi identified cell populations with natural boundaries, while manual gating analysis resulted in abrupt bisecting on some of the 2D plots. Based on the dot plots, the three most difficult-to-resolve gating boundaries seem to be: a) between CD3+CD56+ T (Pop#9) and CD3highCD56+ T cells (Pop#10) (Figure 2C); b) among Naïve CD4+ T and three memory CD4+ T cells (Figure 2D); c) between Tregs (Pop#14) and CD4+ helper T cells (Figure 2E). Dot plots of these cell populations across all 24 samples can be found in Supplementary File 2. Visual examination showed that DAFi successfully identified these difficult-to-resolve cell populations.

Linear regression analysis of the cell population percentages identified by DAFi and CENT for clearly-defined (Figure 2F) and poorly-resolved (Figure 2G) cell populations are highly consistent, with the degree of concordance on the clearly-defined cell populations being higher than that of the poorly-resolved ones. Both *K*-means and DAFi generated highly consistent population percentages with those of CENT (all *p*-values smaller than 0.0001, ranged from *10*^*-4*^ to *10*^*-14*^) for clearly-resolved populations (Figure 2H), indicating that both DAFi and a naive application of *K*-means can identify clearly-defined cell populations successfully. However, *K*-means failed to identify many of the poorly-resolved populations in a consistent way with CENT (Figure 2I), including Tregs, Tcm CD8+ T cells, CD3hiCD56+ T cells, and Temra CD4+ T cells. In contrast, populations percentages of the poorly-resolved populations identified by DAFi are consistent with those derived by CENT (with all *p*-values smaller than 0.01, ranged from *10*^*-2*^ to *10*^*-9*^), indicating the necessity of the recursive filtering and clustering design.

Figures 2J and 2K compare CV (coefficient variability) of population percentages across the 24 samples identified by four different approaches: INDI, CENT, *K*-means, and DAFi. For clearly-defined populations, the four approaches generated highly consistent CV values. However, for poorly-resolved populations, *K*-means and INDI generated very large CV values (Figure 2K), especially for rare cell populations (*e.g.*, Tregs and Temra CD4+ T cells), while DAFi and CENT generated similarly smaller CV values.

### 2.2 Correlation Analysis between Cell Population Proportions with Subject Age and Gender

We extended the single-donor analysis to assess PBMC samples from 132 human subject participants, stained with the same 10-color panel used in Section 2.1. Other details about the FCM experiment can be found in the published study [12]. The goal of the assessment was to determine if T cell population frequency determined by DAFi correlated with subject demographics data, including gender and age.

No difference was observed in age distributions among gender groups (Figure 3A) allowing us to mix the subjects from both genders to increase the statistical power in the age-based correlation analysis. We focused on 12 predefined T-cell populations: 11: T-cells; 12: CD4+ T cells; 13: CD8+ T cells; 14: Tregs; 15: Naïve CD4+ T cells; 16: Tcm CD4+ T cells; 17: Tem CD4+ T cells; 18: Temra CD4+ T cells; 19: Naïve CD8+ T cells; 20: Tcm CD8+ T cells; 21: Tem CD8+ T cells; 22: Temra CD8+ T cells. Complete set of percentages of these T cell populations identified by DAFi across the 132 samples can be found in Supplementary File 3. Consistently with our previous analysis using CENT [12], we found that the proportion of Naïve CD4+ T and Naïve CD8+ T cells decreased with subject age (Figure 3B, corrected linear regression *p*-value 3.740E-06 and 9.678E-11, respectively). Figure 3C shows Pearson correlation scores and linear regression *p*-values across all 12 cell populations identified by DAFi and manual gating analysis. Again, the output of DAFi is highly consistent with that of the centralized manual gating analysis.

**Figure 3.**
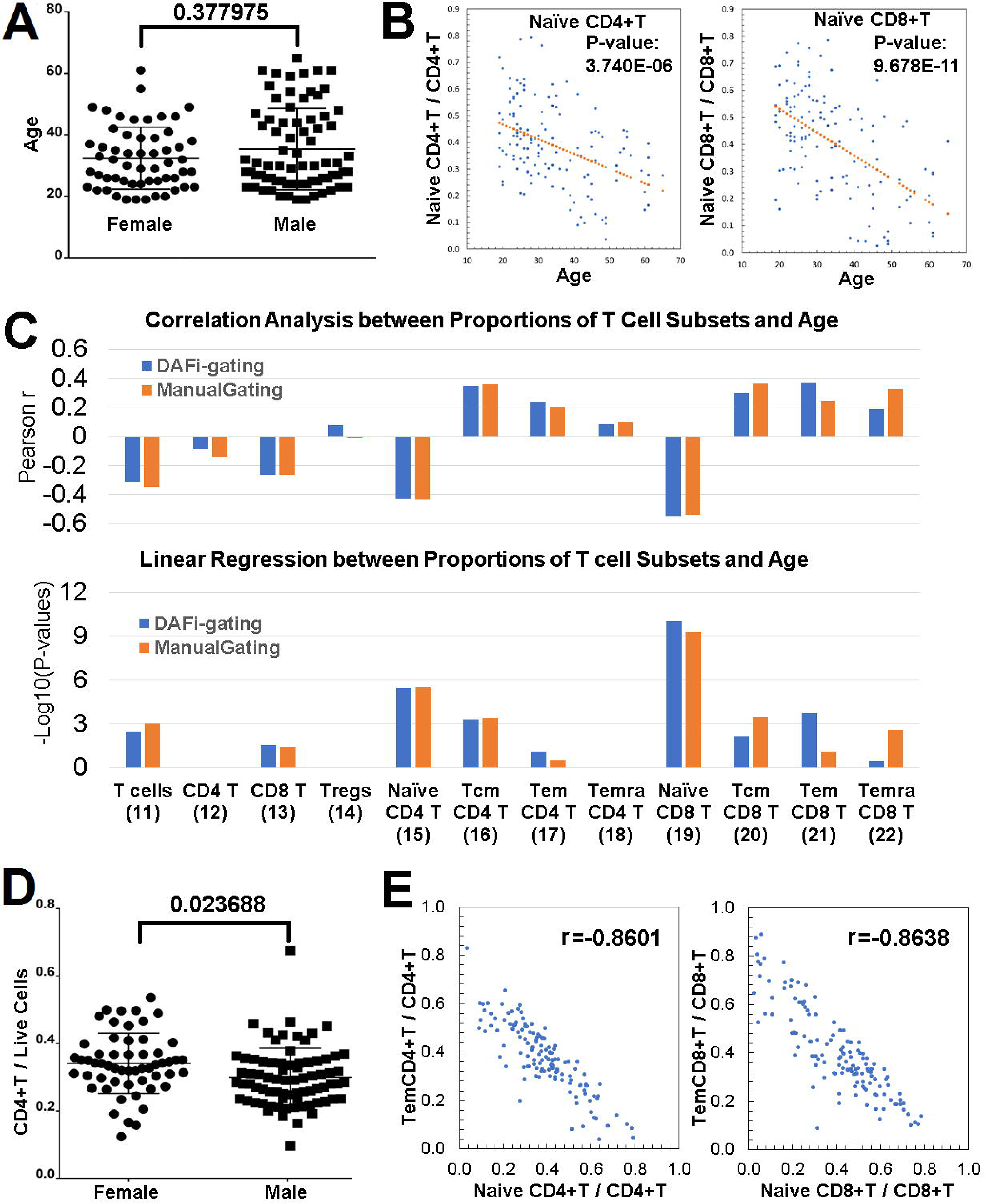
Correlation analysis of DAFi-defined cell population proportions with subject age and gender. A) Age distribution of participants separated by gender. B) Proportions of naïve CD4 T and naïve CD8 T cells (with CD4+ and CD8+ T cells as parents, respectively) versus age with linear regression p-value reported. C) Pearson correlation and linear regression analysis of proportions of T cell subsets with subject age. Parent population definitions of the T-cell subsets can be found in Figure 2A. P-values of x-variable in linear regression analysis were -log10 transformed and multiple comparison corrected by Bonferroni correction. D) Proportion of CD4+ T cells in female and male participants. E) Correlation between the proportions of effector memory T cells versus naïve T cells.

In the gender-based correlation analysis, DAFi identified that the proportion of the CD4+ T cell population seems to be significantly different between the female and male (corrected *p*-value 0.023688, Figure 3D). In our previous analysis [12], we were not able to identify this correlation with a significant *p*-value using manual gating, although the average CD4+ T cell proportion was higher in the female group. A number of previous studies have reported increases in CD4+ T cells in females [4, 17, 20, 37, 39, 40, 41, 42, 43]. Most recently, the 10k Immunomes Project based on a meta-analysis of 578 subjects in the ImmPort Database reported the percentages of CD4+ T cells are significantly elevated in women as compared to men (http://www.biorxiv.org/content/early/2017/08/25/180489).

We also studied the pairwise correlation between population proportions of different T cell subsets. Figure 3E shows the distributions of the population proportion values from the two most significant Pearson correlation scores, both of which are negative. One is between Naïve CD4+ T cells and Tem CD4+ T cells (*r = −0.8601*) and the other is between Naïve CD8+ T cells and Tem CD8+ T cells (*r = − 0.8638*). This finding is consistent with the age-based analytics showing that the number of memory T cells increased with age while the number of naïve T cells decreased with age. For both Tem CD4+ and Tem CD8+ T cell populations DAFi identified a stronger association of their proportion increases with age than manual gating analysis (Figure 3C).

### 2.3 Identification of Known and Novel Cell-Based Biomarkers for Latent Tuberculosis Infection

We also assessed the capability of DAFi for identifying cell populations that have not been defined in the manual gating strategy using a dataset consisting of 12 PBMC samples from 6 latently tuberculosis infected (LTBI) human subjects and 6 6 *Mycobacterium tuberculosis* (*Mtb*) uninfected control (healthy control; HC) subjects used in a previous study [6]. We divided the sequence of manual gating steps into two stages: prefiltering to identify the CD4+ T cell population (the first row of Figure 4A), and unsupervised clustering (FLOCK [33]) to identify cell subsets within the CD4+ T cell population (Figure 4B) associated with subject phenotypes. We noticed that the manual gating strategy after the CD4+ T cells gate was focused on poorly-resolved cell populations with relatively arbitrary gating boundaries on smeared data dimensions including CD25, CCR6, CXCR3, and CCR4 (Figure 4B). Applying a data clustering method that can utilize multiple data dimensions simultaneously to substitute the manual gating strategy starting from CD4+ T cells can be expected to generate more accurate results or identify novel cell-based biomarkers. Complete set of dot plots for data prefiltering by DAFi across all the 12 samples can be found in Supplementary File 4.

**Figure 4.**
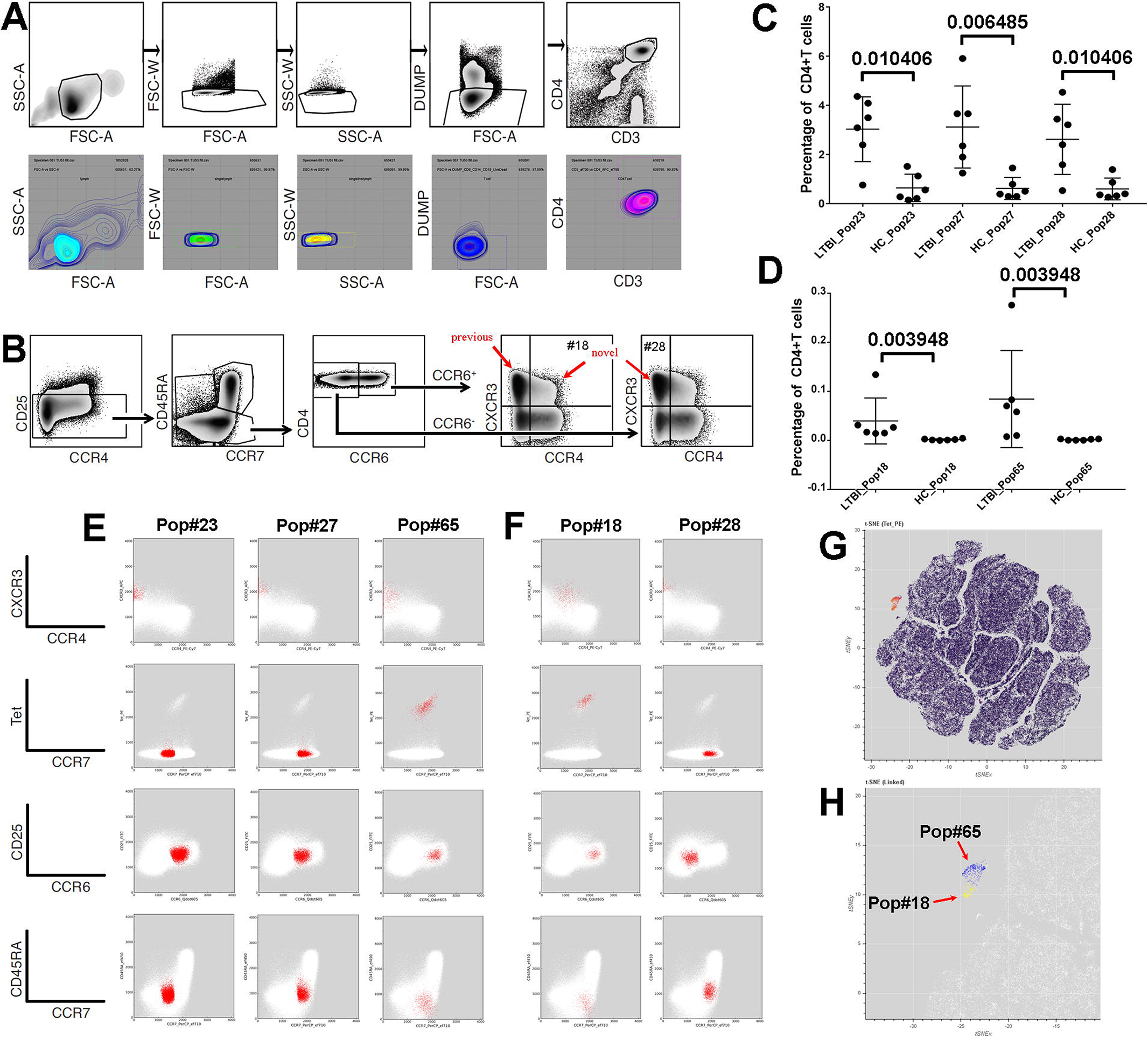
Identification of known and novel cell biomarkers for LTBI using FLOCK-based clustering of DAFi filtered populations. A) Upper: manual gating strategy for identifying CD4+ T cells. The gating path sequentially identifies lymphocytes (FSC-A vs. SSC-A), singlet lymphocytes based on FSC-A/W, singlet lymphocytes based on SSC-A/W, live CD8- T lymphocytes (the DUMP channel includes CD8/CD14/CD19/LiveDead), and CD3+CD4+ T lymphocytes. Lower: CD4+ T cell population identified by DAFi highlighted on corresponding 2D dot plots with density contour lines showing the natural data distribution. A hyper-polygon based on gating boundaries on FSC-A/W, SSC-A/W, DUMP, CD3, and CD4 was used to identify FLOCK data clusters within the CD4+ T cell population. B) Manual gating strategy for identifying subset populations from the CD4+ T cells, based on CD25, CCR7, CD45RA, CCR4, CCR6, and CXCR3 expression. Note that tetramer staining was not used in the manual gating analysis, and different memory T cell regions were not separated based on CCR7 vs. CD45RA. C- D) Percentages of the five most significant DAFi-identified cell subsets (CD4+ T cell population as parent) that differed between LTBI and HC and their corresponding p-values in Wilcoxon rank sum test. Mean and standard deviation of the percentage values of each cell population are shown with the individual values. When N=12, 0.003948 is the best possible *p*-value with the rank sum test when there is no overlap between the ranking of the two groups. E) The three DAFi-identified cell subsets that differ between LTBI and HC in the known CD25-CCR6+CCR4-CXCR3+ region. Events are highlighted in red and shown on different 2D dot plots. The three subsets (Pop#23: Tet-CCR7-CD45RA-, Pop#27: Tet- CCR7+CD45RA-, and Pop#65: Tet+CCR7+CD45RA-) differ from each other based on tetramer and CCR7. F) The two DAFi-identified cell subsets that were not reported in the previous publication with their events highlighted in red on 2D plots of different markers. Both are very rare (average < 0.1% of CD4+ T cells). Pop#18 is Tet+CD25-CCR6+CCR4dimCXCR3+ while Pop#28 is Tet-CD25-CCR6- CCR4-CXCR3+. G) tSNE map of the filtered data. FLOCK clusters of CD4+ T cells (generated without scatter/DUMP/CD3/CD4 values as input) are color-coded based on expression level of tetramer to highlight the tetramer+ population in the mid-upper left region. H) Zoomed-in tSNE map shows that the “island” of the tetramer+ population consists of two separated regions, corresponding to the Pop#18 (highlighted in yellow) and the Pop#65 (highlighted in blue).

DAFi using FLOCK identified 101 cell populations from the 12 samples (percentages of populations can be found in Supplementary File 5). The population percentages were then associated with the subject phenotype using non-parametric Wilcoxon rank sum test with a null hypothesis that there is no difference between the LTBI and HC group. Figure 4C-D shows the top 5 cell populations with most significant *p*-values against the null hypothesis. Distributions of the percentages of three relatively abundant populations (Pop#23, 27, 28) are shown in Figure 4C, with the two rare ones (Pop#18, 65) in Figure 4D. Due to the limited number of subjects, the best possible *p*-value in the rank sum test is 0.003948 when there is no overlapping between the two groups in the ranks of their data objects (Figure 4D).

2D dot plots of the top 5 significant cell populations are shown in Figures 4E-F. In the previous publication [6], a single cell population was identified by manual gating analysis in the CD25- CCR6+CCR4-CXCR3+ region that significantly differed between LTBI and HC in frequency (Figure 4B, corrected *p*-value < 0.01). In contrast, FLOCK identified three subsets within the same region: Pop#23, 27, and 65 (Figure 4E), which differ from each other based on CCR7 and peptide-MHC tetramer staining. DAFi not only identified the known cell-based biomarker but also elucidated the composition of the CD25-CCR6+CCR4-CXCR3+ cell population containing the vast majority of *Mtb*-specific cells. Two DAFi-identified cell populations that were ignored in the original manual gating analysis are: Pop#18 and #28 (Figure 4F), which differ in CCR4, CCR6, and tetramer staining. Their corresponding positions in the predefined cell type hierarchy are indicated with the red arrows “novel” in Figure 4B. Figure 4G shows a tSNE map [5] of the CD4+ T cells of the same LTBI sample used in Figure 4E-F, color-coded based on the tetramer staining levels of the DAFi-identified cell populations. Separation of the tetramer+ cells from the other cells on the tSNE map indicates that two very rare Pops 18 and 65 are indeed distinct (Figure 4H).

### 2.4 Quantification of Human Immune Response to Influenza and Pneumococcal Vaccination

Finally, we applied DAFi to identify plasmablasts/plasma cells to measure human immune responses to vaccination. The FCM dataset used, SDY180 [29], was downloaded from the ImmPort database (www.immport.org). 36 human subjects were enrolled into three immunization arms for FCM experiments: Fluzone (2009-2010 seasonal influenza vaccine, N=12), Pneumovax23 (23-valent pneumococcal vaccine, N=12), and Saline (N=12). PBMC samples were collected at 10 different time points: Day-7, 0 (vaccination day), 0.5, 1, 3, 7, 10, 14, 21, and 28. FCM data files used in our data analysis (306 FCS files in total) were acquired using an 8-color reagent panel focused on identification of the plasmablasts/plasma cells and other types of B-cells: FSC-A, SSC-A, FITC-A_IgD, Pacific-Orange- A_CD45, APC-A_CD138, APC-Cy7-A_CD27, PE-A_CD24, PE-Texas-Red-A_CD19, PE-Cy5-A_CD20, and PE-Cy7-A_CD38. Manual gating analysis was used to identify the cellular composition at the 10 different time points before and after vaccination, which revealed a peak in plasmablast frequencies at Day 7 post vaccination for both vaccines [29].

We reanalyzed the B-cell phenotyping FCM data of SDY180 using DAFi. Dot plots of CD19+ B cells (Figure 5A, blue) and Plasmablasts (Figure 5B, magenta) identified by DAFi are shown with their defining rectangle boundaries. Note that events outside the 2D rectangles may still belong to the cell population as long as the centroid of their data cluster is within the hyper-rectangle. Similarly, an event inside the 2D rectangle may not be assigned to the cell population if its cluster centroid is outside the hyper-rectangle. In Day 7 samples only, the IgD-CD27high plasmablasts can be clearly seen. The clear peaks on Day 7 post both Fluzone and Pneumovax23 vaccinations (Figure 5C) confirmed the finding reported previously [29]. We applied 0-1 min-max normalization to both results (medians of the population percentages across different samples on the same day were used) so that the time-series patterns identified by both approaches could be compared. The time-series pattern identified by DAFi is a close match with that by manual gating analysis (Figure 5D), with a peak on Day 7 for both Fluzone and Pneumovax 23 groups. The second peak post vaccination in the Fluzone group is on Day 14, which also seems a close match between the two approaches. The baseline identified by DAFi seems smoother than that of manual gating analysis in all three groups. 11-fold and 47-fold increase were reported in the previous publication in the absolute numbers of plasmablasts following vaccinations of Fluzone and Pneumovax23, respectively [29]. When comparing the median percentage values of DAFi-identified plasmablasts on Day 7 with the baseline (Day -7), we achieved 16-fold and 43-fold increase post Fluzone and Pneumovax23 vaccinations, respectively, a close match to the manual gating analysis result reported previously.

**Figure 5.**
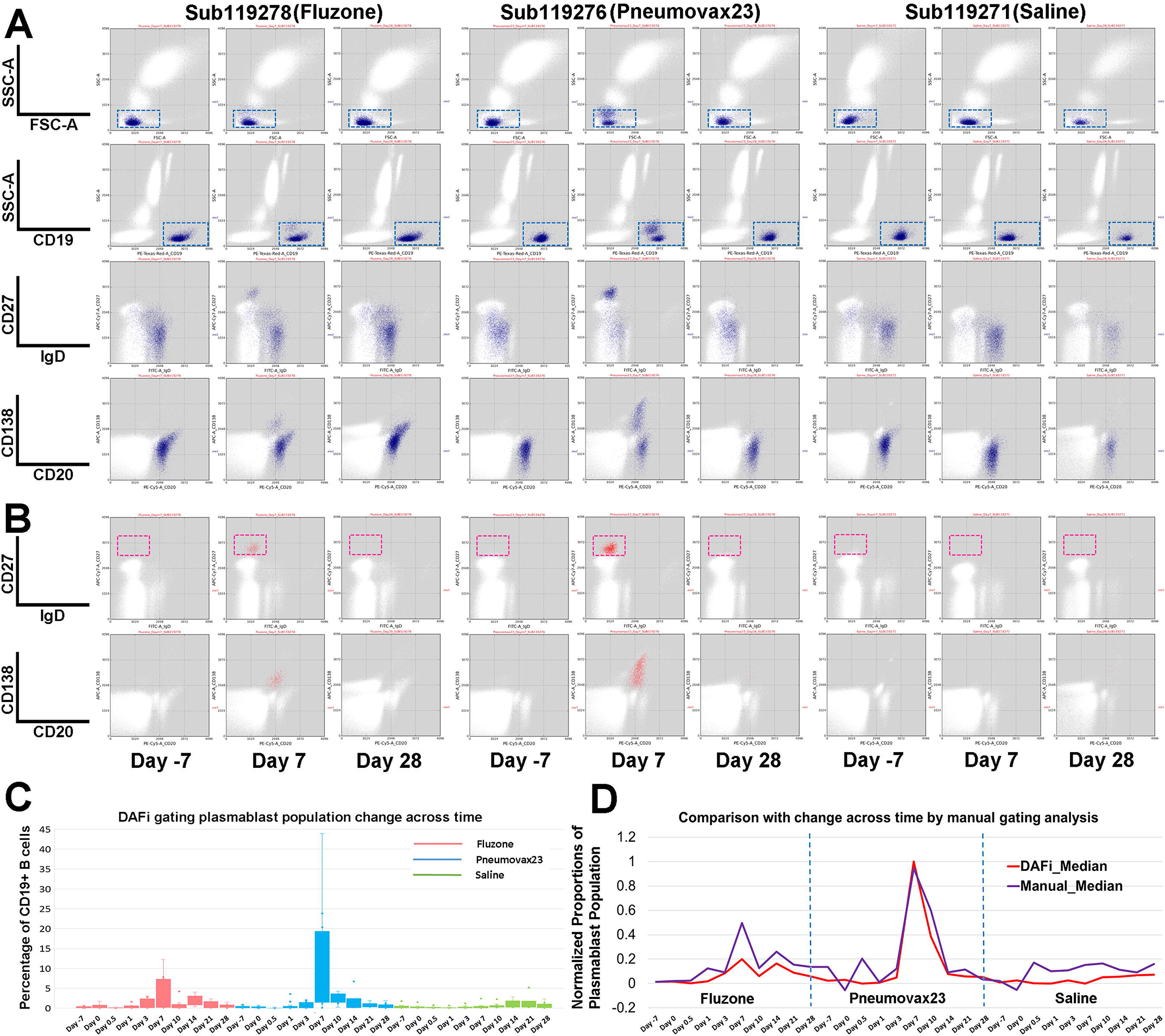
Quantification of human immune response to influenza and pneumococcal vaccination using DAFi. From left to right under each vaccine/saline treatment are three selected time points from one individual in each treatment group: 7 days before the treatment (Day -7), and Day 7 and Day 28 after treatment. A) CD19+ B cells were identified by DAFi using the 2D rectangular gates in FSC/SSC-A and CD19/SSC-A plots illustrated in the first two rows. The two following rows show the B-cell events (colored in blue) on IgD vs CD27 and CD20 vs CD138 dot pots. B) Plasmablast cells identified by DAFi from the CD19+ B cell population. The plasmablasts, defined as IgD-CD27high, are shown in the red box. C) Percentage of plasmablast cells (with CD19+B cell as parent) identified across times and treatment groups by DAFi in box plots. D) Normalized proportions of the plasmablast population (with CD19+ B cell as parent) identified by DAFi and manual gating analysis across times and treatment groups.

## 3. Methods

Design of DAFi consists of four major steps (Figure 1A): unsupervised data clustering, encoding predefined gating boundaries, merging of data clusters based on predefined gating strategy and boundaries, and output for recursive filtering and population statistics. At each gating step, DAFi supports four different gating options/modes to identify the individual cell populations: clustering, bisecting, slope-based bisecting, and reversed filtering. Clustering is the default mode in DAFi. Because of the lack of gold standard in defining and assessing cell populations, we keep the bisecting as an option in DAFi to support the exact recapitulation of manual gating analysis. In some cases, the user prefers to do the gating in a reversed way, *i.e.*, keeping the cells outside of the gate and removing those inside (*e.g.*, the second step in Figure 4B, the identification of memory T cells based on CCR7 vs CD45RA can be achieved by drawing a reversed gate around the naïve T cells in the double positive region), which is supported by the reversed filtering mode in DAFi.

We have implemented two existing data clustering methods: *K*-means and FLOCK to be used with DAFi. We named them DAFi-filtering and DAFi-gating:

DAFi-filtering (for data prefiltering and identification of predefined cell populations):

- Step 1 FCS file preprocessing: each FCS file converted to a data matrix for clustering analysis.
- Step 2 Generation of DAFi configuration file based on manual gating strategy.
- Step 3 Apply *K*-means data clustering to identify data clusters in each input file.
- Step 4 Merge data clusters whose centroids are within the hyper-rectangle formed by gating boundaries; output the merged data as the input file for next-run clustering analysis.
- Step 5 Repeat Steps 3 and 4 until all predefined cell populations of interest are identified.
- Step 6 Output the dot plots and statistics of the identified cell populations together with their names and phenotypes.

DAFi-gating replaces the *K*-means clustering in DAFi-filtering with the FLOCK clustering method that can identify the undefined cell populations:

- Step 1 Run DAFi-filtering with the FLOCK clustering method to identify the predefined cell population that needs to be explored for undefined cell subsets.
- Step 2 Normalize the filtered data across samples and merge the normalized events across samples into a single data file.
- Step 3 Apply FLOCK to the data file to identify data clusters in a fully unsupervised way, and map the clustering results back to individual samples.
- Step 4 Output the dot plots and statistics of the identified cell populations for manual annotation of their names and phenotypes.

### 3.1 FCS file preprocessing

FCSTrans [34] was used to convert and transform (logicle transformation [30]) the binary FCS files generated in all the FCM experiments used in this paper into data matrices for computational processing and analysis.

### 3.2 Converting manual gating strategies into configuration file for DAFi-filtering

Different gating software (FlowJo, FCSExpress, FACS Diva *etc.*) and their different versions use different ways to record the gating boundaries in different formats. Coordinate values of gating boundaries on FCM data from one data transformation cannot be directly applied to gate the data generated by a different transformation or with a different set of transformation parameters. Based on the design of DAFi, we only need to simulate the gating boundaries by drawing rectangles. The Results Section shows that even without using the exact gating boundaries DAFi still achieved highly consistent results with expert manual gating analysis across a variety of experiments and cell populations of interest.

Table 1 illustrates an example configuration file used in DAFi-filtering. The *RecursiveParent* specifies whether this cell population will be used in downstream recursive filtering and clustering. By default, a cell population will be identified from its direct parent in clustering mode, while it can also be identified from its grandparent population or even from the input FCM sample when the user chooses to skip the intermediate gating steps. The former way is recursive clustering while the latter way is not recursive with a hyper-polygon as the constraint, depending on user preference.

**Table 1.**
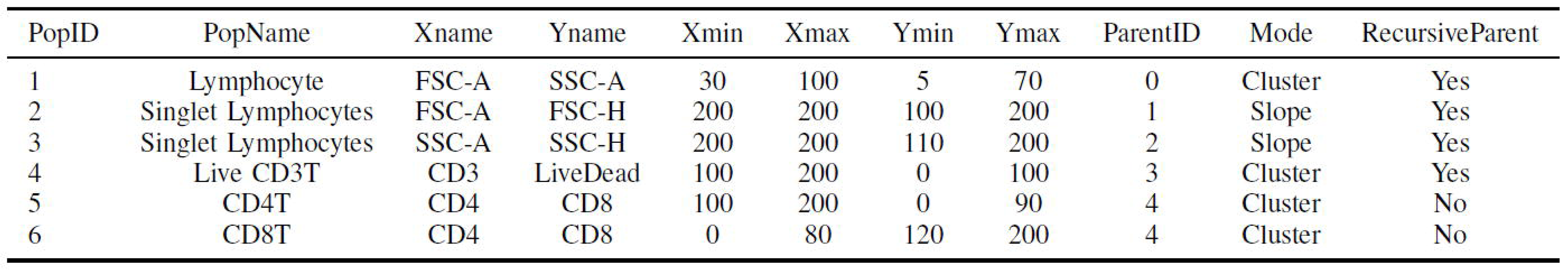
An Example Configuration Table Used in DAFi to Specify Gating Boundary Coordinates and Hierarchical Relationships. Each row corresponds to a cell population defined by the user with Population ID (*PopID*) and Population Name (*PopName*). The two markers used to define the gate were *Xname* and *Yname*. Different data ranges across instruments and experiments are 0-1 min-max normalized into 0-200. DAFi configuration simplified the shapes of the gate by using rectangle gates only. A rectangle gate on a 2D plot is defined by four values: *Xmin*, *Xmax*, *Ymin*, and *Ymax*. For one-dimensional gate on X-axis only, the *Yname* can be an arbitrary dimension and *Ymin/Ymax* can be voided by putting 0/200. The parent of each cell population is specified by the ID of its parent population (*ParentID*), with the *Mode* specifying the way of identifying the cell population from its parent (cluster, slope, bisecting, and reversed).

| PopID | PopName | Xname | Yname | Xmin | Xmax | Ymin | Ymax | ParentID | Mode | RecursiveParent |
| --- | --- | --- | --- | --- | --- | --- | --- | --- | --- | --- |
| 1 | Lymphocyte | FSC-A | SSC-A | 30 | 100 | 5 | 70 | 0 | Cluster | Yes |
| 2 | Singlet Lymphocytes | FSC-A | FSC-H | 200 | 200 | 100 | 200 | 1 | Slope | Yes |
| 3 | Singlet Lymphocytes | SSC-A | SSC-H | 200 | 200 | 110 | 200 | 2 | Slope | Yes |
| 4 | Live CD3T | CD3 | LiveDead | 100 | 200 | 0 | 100 | 3 | Cluster | Yes |
| 5 | CD4T | CD4 | CD8 | 100 | 200 | 0 | 90 | 4 | Cluster | No |
| 6 | CD8T | CD4 | CD8 | 0 | 80 | 120 | 200 | 4 | Cluster | No |

### 3.3 Impact of *K* of *K*-means and size of gating boundaries on DAFi-filtering

While *K*-means is easy to implement and use, one challenge is to set the value of *K*. We have experimented different values of *K* from 100-600 using the LTBI dataset in Section 2.3. Supplementary File 6 shows the *F*-measure values for each of the 5 cell populations comparing between the bisecting (*i.e.*, manual gating analysis) and the clustering mode of DAFi. The box plot shows that the average F1 scores across the 12 samples are larger than 0.95 for all of *K*=100 to 600. The variation of the F1 scores across samples is also small. The larger number of *K*, the closer the result is to the bisecting (the F1 scores seems the largest for *K*=600). By default, we set *K*=500.

We used the same LTBI dataset in Section 2.3 and tested three sizes of gating boundaries for identifying the 5 cell populations across the 12 samples: a) normal: the same size as the bisecting boundaries; b) small: 10% smaller on each dimension than the bisecting boundaries, and c) large: 10% larger on each dimension than the bisecting boundaries. We calculated the precision, recall, and F1 scores of comparing results of DAFi-filtering using these three sets of rectangles against the bisecting results (Supplementary File 7). All the F1 scores as well as precisions and recalls are very high, while using a small rectangle seems increasing the precision but reducing the recall, compared with using a large rectangle. The variation of the F1 scores across the 12 samples is also small without being affected by the slight change of the size of the rectangle gate. For the downstream cell populations (e.g., Pop5: CD3+CD4+ live T lymphocytes), the accumulated change in F1 scores from the previous DAFi-filtering steps is not obvious either.

### 3.4 DAFi-gating: prefiltering and identification of undefined cell populations using FLOCK

For generating the results in Section 2.3, after prefiltering, the remaining events across the 12 samples were first normalized and then merged together. The cross-sample normalization (*a.k.a.*, sample alignment) was done using the GaussianNorm approach [19]. Supplementary File 8 illustrates the application of the GaussianNorm method to normalize the individual data dimensions CCR6 and CD45RA. Only data dimensions needed in the unsupervised FLOCK analysis are kept, resulting in a 7- dimensional data matrix (CD25, CXCR3, CCR4, CCR6, CCR7, CD45RA, and Tetramer), from which FLOCK was applied to identify the 101 cell subsets (*number of bins* = 12 and *density threshold* = 3). The population membership of the events in the merged file is then mapped back to the individual samples for cross-sample statistics and comparisons. Phenotypes of the FLOCK-identified populations were visually examined and manually annotated under the predefined cell type hierarchy (*i.e.*, CD3+CD4+ live T lymphocytes).

## 4. Discussions

The most significant design of DAFi is that it implements the recursive data filtering and clustering along the user-defined manual gating hierarchy, which improves the interpretability of the generated data clusters, including identifying consistent numbers of cell populations across samples and generating user- familiar phenotype definitions of the cell populations. DAFi does not aim to recapitulate the manual gating analysis. Although the manual gating strategy is used, the results of DAFi are data-driven based on the results by unsupervised clustering methods. Both the predefined and novel cell populations identified by DAFi are managed under the same cell type hierarchy for knowledge integration. DAFi can work with different data clustering methods for generating comparable and interpretable results. Because of these characteristics, DAFi will help accelerate the adoption of computational methods by experimental scientists.

The idea of incorporating user inputs into FCM data analysis is not new. Existing approaches such as SPADE [36] and SWIFT [28] require manual operation at the end of the data clustering to group or partition the data clusters into cell populations. Approaches like viSNE [5] and SPADE plot the single cell data in a graph or a transformed space for providing a 2D overview of the high-dimensional data, which can be difficult to interpret or operate on (*e.g.*, grouping the nodes in a SPADE tree into a cell population can be error-prone without checking the events of the nodes on the original 2D plots; a viSNE map is on the tSNE-transformed data space whose dimensions have no biological meaning). In contrast, results of DAFi based on manual gating strategy are much easier to validate and interpret. The use of the manual gating strategy guarantees the consistency of DAFi with manual gating analysis. Supplementary File 9 shows that DAFi outperforms the best unsupervised method on identifying the 4 major cell populations in the FlowCAP-I GvHD dataset in F-measure.

Sample quality control (QC) and cross-sample normalization are important components in any computational pipeline of FCM data analysis. They may impact the results of DAFi. Slight data shifting is not a problem for DAFi. When a data cluster is slightly shifted outside the gating boundaries, its centroid remains within and its events outside of the boundaries will not be lost. However, if there is huge cross-sample variance, currently we need to manually adjust the gates used in DAFi for each group of samples. One solution is to integrate DAFi into a pipeline with components of QC and cross-sample normalization.

Though DAFi was shown to be able to address the existing challenges faced by computational methods, there continue to be improvements needed in the future including eliminating the requirement for a user-provided gating example, which in some cases may be unavailable. For example, one idea is to use flowDensity [27] with DAFi to estimate the boundary coordinates based on 2D data distributions instead of relying on predefined gating boundaries. There are also computational methods being developed to identify the optimal gating path for a given set of cell population phenotypes. The Cell Ontology (CL) [7] can provide a standardized cell type hierarchy to support meta-analysis across different FCM experiments. Development of a graphical user interface for allowing the data analyst to create different gating sequences, connect with FlowJo workspace files, compare result statistics, and integrate with other data filtering and clustering methods will help improve the usability of DAFi.

We implemented and benchmarked the performance of DAFi on the Comet cluster at the San Diego Supercomputer Center. Through parallel computing, DAFi processing and analysis of the 306 files in the ImmPort SDY 180 was completed in about 30 minutes using a single compute node with 24 CPU cores. We are collaborating with FlowJo to release DAFi as a plug-in tool. We are also integrating DAFi into the FlowGate cyberinfrastructure [35], and implementing it on a JupyterHub server for interactive auto-gating analytics of FCM data.

## 5. Conclusions

The advancement of FCM data with increased dimensionality brings in challenges in data analytics, but also provides possibilities for the identification of novel cell-based biomarkers based on measurements on additional combinations of markers. The large number of measured characteristics also provide information to accurately define a cell population, which is facilitated by the development and use of computational methods for automated identification of cell populations. How to integrate human intelligence on pattern recognition with the power of computation to identify cell populations from high- dimensional FCM data robustly and interpretably is a challenge that has not been sufficiently addressed. In this paper, we propose a new computational method and framework - DAFi. Datasets from four different study settings were used to evaluate the performance of DAFi, demonstrating DAFi’s characteristics in

- Generation of consistent cell type specific statistical measurements with expert centralized manual gating analysis;
- Identification of natural shapes of both major and rare cell populations;
- Identification of both clearly-defined and poorly-resolved cell populations, and
- Easy interpretation and management of the identified cell populations using user-defined manual gating strategy.

## Acknowledgments

La Jolla Institute for Allergy and Immunology: Veronique Schulten, Jason Greenbaum; Human Longevity Inc.: Rick Stanton; University of Rochester: David Topham, David Roumanes, Edward Walsh, Gloria Pryhuber, Nathan Laniewski, Kristin Scheible, Jeanne Holden-Wiltse; FlowJo LLC.: Josef Spidlen; San Diego Supercomputer Center: Robert Sinkovits.

**Author Contributions:**

DAFi method design: YQ and RHS; DAFi method implementation: YQ, IC, and AJL; Computational processing and analysis of FCM data: AJL, IC, YQ, and JB; Flow cytometry data acquisition and manual gating analysis: JB, CLA, and DW; Immunology use cases and result interpretation: RHS, BP, and AS; Manuscript preparation: YQ, AJL, and RHS. All authors helped with revision of the manuscript. All authors read and approved the final manuscript.

**Availability of Data and Software:**

The FCM datasets used in this manuscript have been submitted to FlowRepository under accessions:

FR-FCM-ZYBS, FR-FCM-ZYBT, FR-FCM-ZYBU

The HIPC study data used in this manuscript is also publicly available at the ImmPort database (SDY820); the ImmPort SDY 180 dataset is publicly accessible at ImmPort (https://www.immport.org) and can be downloaded at: https://aspera-immport.niaid.nih.gov:9443/browser?path=SDY180

## Supplementary Materials

Supplementary File 1: Definitions of 22 cell populations using a 10-color T-cell FCM reagent panel.

Supplementary File 2: Different T cell subsets identified by DAFi across 24 repeat runs of PBMC from a control donation, including clearly defined and poorly resolved ones.

Supplementary File 3: Proportions of the 22 predefined cell populations identified by DAFi across 132 subjects for correlation with age and gender

Supplementary File 4: Results of DAFi prefiltering by applying FLOCK clustering method across 12 samples in LTBI study. From left to right: lymphocytes, singlet lymphocytes, live singlet lymphocytes, and CD3+CD4+ live T lymphocytes.

Supplementary File 5: Population percentages of the 101 cell populations identified by FLOCK (with CD3+CD4+live T lymphocytes as parent) and their rank sum test *p*-values between LTBI and HC groups.

Supplementary File 6: F1 scores of using different *K* in *K*-means clustering for DAFi-filtering across the 12 samples in LTBI study for each cell population compared with bisecting analysis.

Supplementary File 7: Precision, recall, and F1 scores when using different sizes of gating boundaries in DAFi-filtering across the 12 samples in LTBI study for each cell population compared with bisecting analysis.

Supplementary File 8: Results shown in histograms before and after applying GaussianNorm method to normalize CCR6 and CD45RA channels across the 12 samples in LTBI study.

Supplementary File 9: Comparison of dot plots and F1 score between manual gating analysis in FlowCAP-I GvHD dataset and the DAFi result.

